# Relationships between individual differences in dual process and electrophysiological signatures of familiarity and recollection during retrieval

**DOI:** 10.1101/2021.09.15.460509

**Authors:** Halle R. Dimsdale-Zucker, Karina Maciejewska, Kamin Kim, Andrew P. Yonelinas, Charan Ranganath

## Abstract

Our everyday memories can vary in terms of accuracy and phenomenology. According to one theoretical account, these differences hinge on whether the memories contain information about both an item itself as well as associated details (remember) versus those that are devoid of these associated contextual details (familiar). This distinction has been supported by computational modeling of behavior, studies in patients, and neuroimaging work including differences both in electrophysiological and functional magnetic resonance imaging. At present, however, little evidence has emerged to suggest that neurophysiological measures track individual differences in estimates of recollection and familiarity. Here, we conducted electrophysiological recordings of brain activity during a recognition memory task designed to differentiate between behavioral indices of recollection and familiarity. Non-parametric cluster-based permutation analyses revealed associations between electrophysiological signatures of familiarity and recollection with their respective behavioral estimates. These results support the idea that recollection and familiarity are distinct phenomena and is the first, to our knowledge, to identify distinct electrophysiological signatures that track individual differences in these processes.

## Introduction

We have all had the experience of running into someone we know walking down the street. Sometimes just seeing that person can help us recollect their name and where and when we last encountered them. At other times, we might feel confident that we have met that person because the face seems so familiar, even if we are otherwise unable to recover any details about that person. A long history of memory research has suggested that recollection and familiarity vary in terms of retrieved information (i.e., item vs. context information), vividness (Cooper and Ritchey, 2019; Jacoby and Dallas, 1981; Roediger and Blaxton, 1987; Tulving, 2002; Woroch and Gonsalves, 2010), and subjective experience (Leynes and Nagovsky, 2016; Souchay et al., 2013). Moreover, studies using electroencephalography (EEG; Addante et al., 2012b; Curran, 2002; Diana et al., 2011; Duarte et al., 2004; Düzel et al., 1997; Leynes and Phillips, 2008; Rugg and Curran, 2007; Tsivilis et al., 2001; Wilding et al., 1995; Woroch and Gonsalves, 2010), magnetoencephalographic (Evans and Wilding, 2012), studies of patients with brain damage (Addante et al., 2012a; Aggleton et al., 2005; Aly et al., 2011; Bowles et al., 2010; Duarte et al., 2005, 2004; Wang et al., 2014, 2014), and functional magnetic resonance imaging (fMRI; Diana et al., 2012, 2007; Eldridge et al., 2000; Vilberg et al., 2006; Vilberg and Rugg, 2008; Yonelinas et al., 2005) are consistent with the idea that recollection and familiarity depend on different neural substrates. Accordingly, many theories (Jacoby, 1991; Mandler, 1980; Tulving, 1985; Yonelinas, 1999) and computational models (Elfman et al., 2014; Norman and O’Reilly, 2003; Selmeczy and Dobbins, 2014) have proposed that recollection and familiarity are driven by different processes (see Eichenbaum et al., 2007; Ranganath, 2010a, 2010b; Ranganath and Rainer, 2003; Reagh and Ranganath, 2018; Wilding and Ranganath, 2012; Yonelinas, 2002 for extensive reviews of this literature), though this idea remains somewhat controversial (e.g., Wixted, 2007).

One class of models (Park and Donaldson, 2019; Yonelinas, 2002, 1999, 1994; Yonelinas et al., 2010) particularly emphasizes the idea that recollection and familiarity can be distinguished on the type of associated details recovered, as well as phenomenology. According to this view, familiarity is typically associated with strong memory for an item with minimal information about associated contextual details, whereas recollection is accompanied by retrieval of information about the item and about the context in which it was encountered (Addante et al., 2012a; Diana et al., 2008; Eichenbaum et al., 2007; Ranganath, 2010b; Yonelinas, 2002, 1999, 1994; Yonelinas et al., 2010).

Many studies have used recordings of event-related potentials (ERPs) to differentiate between neural correlates of recollection and familiarity based on sensitivity to particular experimental variables, temporal dynamics, and scalp topography (Addante et al., 2012b; Bridger et al., 2012; Curran, 2004, 2000; Park and Donaldson, 2019; Rugg et al., 1998a, 1998b; Rugg and Curran, 2007; Wilding and Herron, 2006, 2006; Wilding and Ranganath, 2012), but there is significant controversy over how these results should be interpreted (Hou et al., 2013; Lucas et al., 2010; Nie et al., 2014; Paller et al., 2007; Thakral et al., 2016; Voss et al., 2012; Voss and Paller, 2016, 2006; Yovel and Paller, 2004). Many studies have reported ERP differences between old and new items (i.e., an ERP “old-new effect”) around 300-600 ms post-stimulus, with a mid-frontal or fronto-central scalp topography (Curran et al., 2006; Friedman and Johnson, 2000; Mecklinger, 2006, 2000; Rhodes and Donaldson, 2007; Tsivilis et al., 2001). The topography and latency of this “mid-frontal old-new effect”(Rugg et al., 1998a) resembles the N400 ERP component reported in psycholinguistic studies (Kutas and Hillyard, 1984), although the mid-frontal old-new effect often has a more anterior distribution (Wilding and Ranganath, 2012). The mid-frontal old-new effect has often been contrasted with a late-onsetting old-new effect that is maximal at parietal sites. This latter effect has been called the “parietal old-new effect” or the late positive component (LPC) (Friedman and Johnson, 2000; Olichney et al., 2000; Paller and Kutas, 1992; Smith, 1993; Wilding and Ranganath, 2012), and is more left-lateralized for words and more widespread for pictures and actions (Leynes et al., 2017).

Results from a number of studies have been used to support the idea that the mid-frontal old-new effect may be a neural correlate of familiarity, whereas the parietal old-new effect may be a neural correlate of recollection (Addante et al., 2012b; Friedman and Johnson, 2000; Griffin et al., 2013, 2013; Leynes et al., 2005; Olichney et al., 2000; Paller and Kutas, 1992; Park and Donaldson, 2019; Rhodes and Donaldson, 2007; Rugg et al., 1998b; Speer and Curran, 2007; Wilding and Ranganath, 2012, 2012; Wynn et al., 2020, 2020) for reviews see (Curran et al., 2006; Friedman, 2013; Friedman and Johnson, 2000; Rugg and Curran, 2007), although results from other studies have argued against this idea (Bridger et al., 2012; Greve et al., 2007; Kelley and Wixted, 2001; Leynes et al., 2017; Paller et al., 2007; Voss and Federmeier, 2011; Voss and Paller, 2006; Yovel and Paller, 2004). One complication in interpreting studies differentiating between the mid-frontal and parietal old-new effects is that the exact timing and topography of these effects can differ considerably across studies, and it is possible that recollection and familiarity might be differentiated by ERP modulations that do not exhibit the typical characteristics of these two old-new effects (Friedman et al., 2005; Ranganath and Paller, 2000; Tsivilis et al., 2001). Data-driven ERP analysis methods (Maris and Oostenveld, 2007) might offer a way to reconcile these views and identify the extent to which recollection and familiarity can be differentiated.

It is also notable that most ERP studies have focused on old-new effects in group averages, and little is known about whether ERP modulations track individual differences in memory performance. If different ERP measures are predictive of putatively different memory processes, there should be unique correlations between the behavioral and ERP measures of each memory process. Recently, several studies have focused on correlating ERPs with individual differences in recognition memory performance (Amico et al., 2015; Angel et al., 2010; Chen et al., 2014; MacLeod and Donaldson, 2017), but these studies have not revealed a clear picture. Angel et al. (2010) correlated overall recognition memory performance (corrected recognition rate) with the magnitude of the parietal old-new effect, but this study was performed on a small sample (14 participants), and it focused only on the parietal old-new effect within an a priori time widow. More recently, MacLeod and Donaldson (2017) correlated the magnitude of the left parietal old-new effect with recognition performance. Across three tasks, the authors found significant old/new effects in the left parietal ERP (Hit>CR; R-CR > K-CR), but correlations between the ERP and behavioral measures were inconclusive. This may have arisen because, although the total number of participants was high (122), only 20 participants were included in the correlation of R/K effect magnitude with behavioral data. In addition, these analyses focused only on the late left parietal effect, estimated as the mean ERP difference within an a priori time widow, averaged across three parietal electrodes and did not attempt to differentiate correlates of recollection- and familiarity-based recognition. Chen et al. (2014) correlated FN400 (mid-frontal old-new effect) magnitudes with recognition performance in a large sample (64 participants), but this study used only overall recognition discriminability (d′) and response time as behavioral indices of memory performance, which does not distinguish recollection from familiarity. Whereas the above studies focused specifically on previously identified ERP old-new effects, Amico et al. (2015) used data-driven non-parametric analyses to characterize individual differences in overall recognition performance (Hit and FA rates, the sensitivity index d’, the decision criterion c, and the mean RT for Hit trials). However, this study had a relatively small sample size (18 participants) and did not attempt to separately estimate familiarity and recollection.

To summarize, prior studies have focused on relationships between ERP components and recognition memory performance, but no previous study, to our knowledge, has shown a relationship between individual differences in behavioral estimates of recollection and familiarity and the putative ERP correlates of these processes described above. Furthermore, given the known variability in timing and topography in these EEG signatures with different types of stimuli, there is no agreed upon standard definition that can be applied consistently across studies.

The present study seeks to address these limitations by testing the hypothesis that individual differences in recognition memory performance are related to variability in electrophysiology in terms of two recognition memory processes: familiarity and recollection. In order to test whether individual differences in electrophysiological signatures of familiarity and recollection were related to differences in their dual process estimates, we measured ERPs while participants made recognition memory judgments using the Remember/Know method (Tulving, 1985). To identify ERPs related to individual differences in familiarity and recollection in an unbiased manner, we used a data-driven analysis approach in which ERP differences were correlated with individual differences in dual process estimates of familiarity and recollection using non-parametric cluster-based permutation analysis. If dual process estimates of familiarity and recollection are associated with particular electrophysiological signatures, we should observe correlations between familiarity-related ERPs and familiarity dual process estimates and between recollection-related ERPs and recollection dual process estimates.

## Methods

### Participants

49 participants took part in the study. They were right-handed and had normal or corrected-to-normal visual acuity and no history of neurological or psychological disorders. Participants were excluded due to contamination of mastoid channels (N=1), incidental MRI finding (N=1), not completing the task (N=2), and not fulfilling the inclusion criteria of at least 30 recollection trials (Boudewyn et al., 2018; Cohen, 2014; Luck, 2014, 2005), N=7). Thus, 38 participants (N_female_ = 26, mean age = 25.7±4.2 years) were included in the final analyses. A post-hoc sensitivity analysis of a bivariate normal model correlation using G*Power 3.1 software (Faul et al., 2009, 2007) with a two tailed alpha value of 0.05 showed that a sample size of 38 with a power of 0.8 could detect a medium to large effect size of 0.436 (Cohen, 1988) with a correlation interval of ±0.32. The study was approved by the Institutional Review Board of the University of California at Davis and all participants provided informed consent prior to participation. Participants were compensated $20/hr for their time.

### Stimuli

Study materials came from the Bank of Standardized Stimuli (BOSS) developed by Brodeur and colleagues (Brodeur et al., 2014, 2010). Using the normative data provided by the BOSS creators, we selected the images that were well-agreed on for the name (>29%), category (>29%), object (>2 on a 1-5 scale), and viewpoint (>2 on a 1-5 scale), and rated as familiar (>2 on a 1-5 scale), and simple (<2 on a 1-5 scale for complexity). Duplicate items were removed as well as items in any of the animal (“Crustacean”, “Mammal”, “Reptile”, “Bird”, “Insect”, “Canine”, “Feline”, “Sea mammal”, “Fish”), body part, and war weapon categories. In addition, any potentially emotional, disturbing, or unpleasant images were removed (e.g., syringe, hunting knife) as well as objects that a research assistant deemed hard to recognize (e.g., contact lens) or redundant with another object. Images were re-sized in Adobe Photoshop to 500 x 500 pixels. Objects from this list of 644 remaining objects were then used to generate study and lure lists. Object categories used in the stimuli represent the general makeup of the original BOSS stimuli (see Brodeur et al., 2014, 2010 for details). Original stimuli used in the paradigm (https://osf.io/4s7uy/) as well as the original source code for stimulus presentation (https://github.com/hallez/eetemp_eeg_pub/tree/main/experiment-scripts) are available online.

270 objects were uniquely drawn for each subject such that each participant saw a distinct set of items although, by chance, some objects necessarily overlapped between participants. Of the 270 objects selected for a participant, 180 images were used as study items during the encoding phase and 90 were used as lures during the retrieval phase. 180 study items were divided into five encoding lists of 36 items each. One of four encoding questions (“Would you find this item in a supermarket/convenience store?”, “Would this item fit in a fridge/bathtub?”) was paired with each item. In each of the five encoding lists, each question was paired with 9 different items such that each question was used the same number of times in each encoding list. Lists for object retrieval were constructed by randomly selecting 30 studied and 15 lure items. The proportion of objects from each encoding list (1-5) and encoding question (fridge/bathtub/supermarket/convenience store) was not constrained in the construction of the retrieval lists. This manipulation was used to ensure the participants were paying attention to individual items and was not included in the analyses described in this work. For a retrieval list, the 180 studied items and 90 lure items were randomly intermixed, using the *numpy.random.shuffle* tool in Python (Version 2.7.14). Participants were given a practice phase with four objects. Objects used in the practice phase were drawn from a separate stimulus set of computer-generated images (previously used in (Dimsdale-Zucker et al., 2018) so that they would not be confused with any of the studied items. Four practice objects were randomly selected for each subject from a set of 12 objects.

### Procedure

The experiment comprised of practice, encoding, and retrieval phases. EEG data were collected during the retrieval phase. Each trial during the retrieval phase included object recognition and source memory test components. The present study focuses on the EEG signature that differentiates recollection and familiarity during item recognition as measured with respect to the object recognition judgment.

For practice, a research assistant walked the participant through each phase of the task. The practice for encoding was followed by familiarizing participants with the response scales that were used for the object recognition and source judgments. Participants were told that the difference between familiarity and recollection is that familiarity is feeling like they know or have seen this item but being unable to recall from where. In contrast, recollection is accompanied by memory for both the item itself as well associated details (e.g. an experience or association from when the item was originally studied, the source or context of the item).

This description was supplemented with an example of meeting someone in the grocery store and either feeling like you know them but not having access to their name or how you know them (“familiar”) versus running into an individual and knowing their name or where you’ve encountered them previously (“remember”). New responses were explained as being analogous to the experience of meeting a stranger or a totally novel person. The participants were given practice trials along with feedback from the experimenter and detailed instructions to ensure they understood how to use this response scale.

During the encoding phase, objects and questions (“Would this item fit in a fridge?”, etc.) appeared on the screen for 250 ms. Within each list, all four questions were presented in a random order, an equal number of times across items. The participant made their response to the question during this interval. A short presentation time was used in order to eliminate potential eye movements during encoding. Response key mappings for the yes/no judgments were randomized across participants. Responses were recorded but accuracy was not analyzed as the purpose of this task was simply to orient participants to the objects and their respective sources. Between lists, there was a 30 second break before the text label (e.g. “List 1”) for the next list was shown.

After studying all 180 objects, participants were then setup for EEG recording. Details of the EEG setup can be found in the *EEG acquisition and processing* section. This cap setup served as the delay period (approximately 45-60 min) between the encoding and retrieval phases. During the retrieval phase, participants saw 270 objects (180 old and 90 novel) split across six blocks of 45 trials (30 old and 15 novel objects). Each block began with a 23 second warm up period that walked the subject through getting in a comfortable position, blinking, and preparing to begin responding while allowing the EEG recordings to stabilize. Each trial started with a white central fixation cross on top of a gray background that remained in the foreground of the screen throughout the entire trial. Next, an object appeared and remained on the screen for 700 ms (Figure 1). Participants knew to withhold their response at this time. A capital white “T” (“think cue”) came on screen for 1700 ms. This timing was based on a similar paradigm (Gruber et al., 2008). Again, no responses were made while the think cue remained on the screen to minimize movement-related artifacts in the EEG data in the time window of interest.

**Figure 1.**
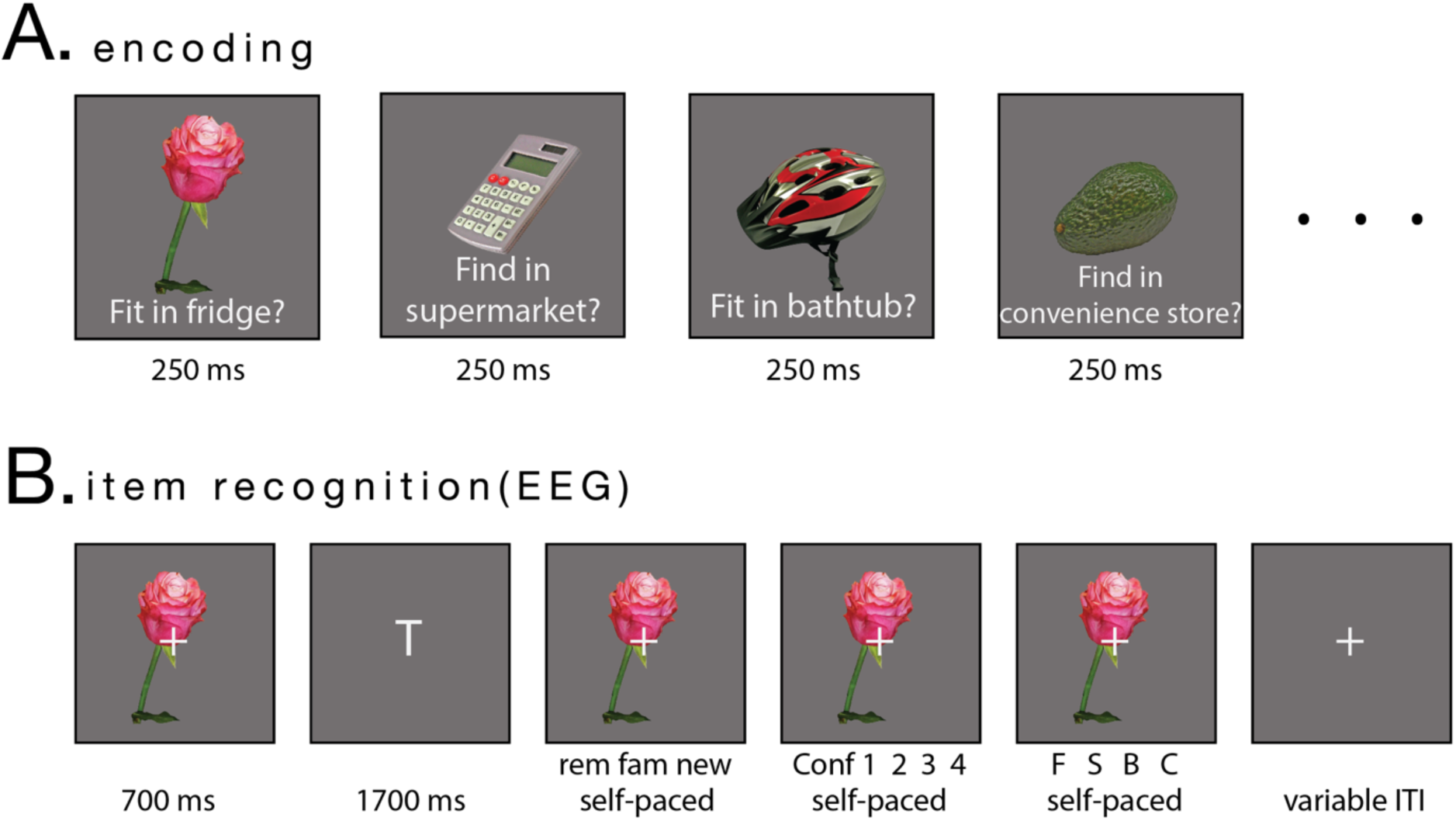
Paradigm design. During the encoding phase (A) participants studied items and answered one of four questions: “Would you find this item in a supermarket/convenience store?”, “Would this item fit in a fridge/bathtub?”. Items were presented on top of a gray background with questions written in white text. During (B) participants saw old and new items, while undergoing EEG recordings. Items were presented on a gray background with a white fixation cross overlaid. Each item was presented briefly (700 ms). Initial item presentation was followed by a think cue (white letter “T”). Following this, participants made their memory response to the item (rem =Remember, fam =Familiar, new =New item), followed by a memory confidence level judgment and a question source memory judgment (“fridge”, “supermarket”, “bathtub”, “convenience store”; abbreviated above in figure as “F S B C”). ERPs were recorded during the item recognition phase. Black outlines are for figure visualization purposes only and did not appear during the task to participants.

Finally, the object reappeared, and the participant was allowed to make a self-paced object recognition judgment (Remember, Feels Familiar, or New; response order counterbalanced). After making their memory judgment, the object remained on the screen but the response scale updated to a confidence judgment (1=highly, 2=moderately, 3=somewhat, 4=not at all, Figure 1). This was followed by a four-option forced-choice question about the item’s original encoding question (again, the order of the four options was randomized for each subject but remained constant throughout the experiment). Trials were separated by 2 second ITIs with a white fixation cross. For EEG timing precision, the timing of all screens was taken as the ceiling of the expected duration multiplied by the monitor’s frame rate.

### Behavioral measures

Behavioral measures of recollection and familiarity were calculated using dual process estimates, which are intended to give an independent measure of these two processes, while accounting for response bias (Yonelinas, 2002). Recollection was estimated as the difference between the Remember hit rate and the Remember false alarm rate (*[count of all remembered trials / count of all old items] – [count of all remembered false alarms / count of all new items]*) and familiarity was estimated as the difference between the Familiar hit rate corrected by the inverse Remember hit rate and Familiar false alarm rate corrected by the inverse Remember false alarm rate (*[Familiar hit rate / (1 – Remember hit rate)] – [Familiar falsealarm rate / (1 – Remember false alarm rate)],*(Yonelinas, 2002). Source judgments to old items were computed to determine whether participants could retrieve information about the question. We expected that accurate retrieval of the orienting task used to encode each item (source memory) should be more likely for items judged as recollected (MacKenzie et al., 2018; Yonelinas, 2002) than for items associated with “familiar” responses.

### EEG acquisition and analysis

EEG data were recorded in a sound attenuated chamber at a rate of 512 Hz using a BioSemi (http://www.biosemi.com) ActiveTwo system with 64 active Ag/AgCl scalp electrodes embedded in an elastic cap in an extended version of the international 10/20 system. Additional electrodes were placed at the left and right mastoids to be used for offline re-referencing. In addition, the electrooculogram (EOG) was recorded with an additional four electrodes – bipolar vertical channels located approximately 1cm above and below the subject’s left eye and horizontal ocular channels located approximately 1 cm lateral to the outer canthus of each eye. The EEG was recorded relative to a common mode sense active electrode near Cz for online referencing. Participants were instructed to blink normally while maintaining focus at the center of the screen and while minimizing muscle tension and any large movements.

EEG data preprocessing and analyses were performed using EEGLab (Delorme and Makeig, 2004), ERPLab (Lopez-Calderon and Luck, 2014), and custom code implemented in MATLAB r2014b (www.mathworks.com) for EEG processing, available in a public github repository (https://github.com/hallez/eetemp_eeg_pub/tree/main). Data intended for ERP analyses were downsampled offline to 128 Hz, re-referenced to the average of the mastoid channel signals, and high-pass filtered at 0.1 Hz (IIR Butterworth filter, half-amplitude cutoff=0.2 Hz, slope=12 dB/octave). The data were separately high-pass filtered for ICA (Winkler et al., 2015) at 1Hz (IIR Butterworth filter, half-amplitude cutoff=1.60 Hz, slope=12 dB/octave), based on Makoto Miyakoshi’s preprocessing pipeline (https://sccn.ucsd.edu/wiki/Makoto's_preprocessing_pipeline). Next, bad channels were detected using the *trimOutlier* function (lower standard deviation threshold=2; upper standard deviation threshold=200) and then epoched (epoch start = −500ms; epoch end = 1000ms). After epoching, outlier epochs were automatically rejected using a thresholded approach implemented with *pop_jointprob* (local channel threshold = 6; global threshold = 2). Independent component analysis (ICA) was performed with the *runica* algorithm (Delorme et al., 2007) holding out mastoid and outlier channels, and the SASICA toolbox (Chaumon et al., 2015)was used to manually review and identify eyeblink-related components for removal from the data. At manual review, any additional epochs or channels that were determined to be outliers were identified for removal or interpolation, respectively. Bad channels were interpolated from the ICA corrected data (mean number of interpolated channels was 1.3±1.9 per block). At this point, all six blocks were merged into a single file. Baseline correction was applied to each trial using a pre stimulus period from 200 ms prior to the onset of the first image in a trial. Average number and standard deviation (with the range provided in parentheses) of trials included in the analysis was: 66±12 (34-83), 54±15 (33-94), and 79±24 (30-127) for Correct Rejection, Familiar, and Remember trial types, respectively. Raw and preprocessed files can be found at https://osf.io/4e3pq/.

Given that ERPs related to long-term memory are known to shift their latency and distribution with different types of stimuli (Addante et al., 2012b; Busch et al., 2004; Taylor, 2002; Yonelinas, 2002), we performed non-parametric cluster-based permutation analysis (Maris and Oostenveld, 2007) implemented in the FieldTrip Matlab toolbox (Oostenveld et al., 2011), which corrects for the multiple comparisons problem (MCP) arising from the fact that the effect of interest (i.e. a difference between experimental conditions) is evaluated at large number of data points, here: (channel, time)-pairs. The approach combines neighboring values that are likely to be correlated (e.g., neighboring time points and/or spatial locations) to reduce the problem of multiple comparisons. Therefore, this method allowed us to compare ERPs between trial types (Familiar, Remember, and Correct Rejection) for each (channel, time) data pair and identify statistically meaningful differences. Under the null hypothesis of exchangeability, assuming averages from Familiar, Remember, and Correct Rejection trials are drawn from the same probability distribution, cluster alpha *p* =0.05 and 0-700 ms time window were used. In order to isolate familiarity and recollection electrophysiological estimates while controlling for other cognitive processes that are not related to memory, ERPs from the following trial types were compared: Familiar as compared to Correct Rejection trials, and Remember as compared to Familiar trials, respectively. Such comparisons are widely used in the studies of old-new effect, including Remember/Know procedures (Duarte et al., 2006, 2004).

The comparisons between each pair of trial types were performed via a two-tailed t-test (alpha = 0.05) using *depsamplesT* function, which clustered samples whose t-value was larger than a priori threshold (*P*=0.05) on the basis of temporal and spatial adjacency. Cluster-level statistics were calculated by taking the sum of the t-values within every cluster and the maximum of the cluster-level statistics was taken. Monte Carlo correction for the MCP with 1000 draws from the permutation distribution was used. Channel neighbors for spatial clustering were found based on the template method, using ‘Biosemi64_neighb’ template.

In order to test our critical question of whether ERP signatures associated with recollection and familiarity are associated with dual process estimates, we computed correlations between these measures. This was also performed using a non-parametric cluster-based permutation test. In this case, *Ft_statfun_correlationT* function and Pearson *r* coefficient were used in order to test if there was a relationship between familiarity and recollection dual process estimates per subject (quantitative independent variable) and their (channel, time) EEG data (dependent variable). The correlations of both difference waveforms (Familiar – Correct Rejection, and Remember – Familiar) with behavioral estimates of familiarity and recollection, as generated from dual process estimates, were tested.

## Results

### Behavioral results

Participants were highly accurate at discriminating studied from unstudied objects (Table 1). To account for response bias, we also computed dual process estimates of familiarity and recollection (Yonelinas, 2002). Mean familiarity and recollection estimates are presented in Table 1. Each studied item was scored according to the item recognition judgment and accuracy for the question type the item had been paired with (Table 1). The accuracy of source memory was significantly above chance for both familiarity (t_37_ = 7.0, *P*< 0.001) and recollection (t_37_ = 10.3, *P*< 0.001), but participants were significantly more likely to correctly retrieve source information for Remember than Familiar trials (t_37_= 6.0, *P*< 0.001). These findings are in agreement with other work showing that accurate source judgments can be made on the basis of both recollection and familiarity (Addante et al., 2012b; Diana et al., 2011, 2008), but that retrieval of contextual details should be more likely when an item is recollected, as compared with familiarity-based recognition (for a comprehensive review, see Yonelinas, 2002).

**Table 1.** Behavioral results of object recognition and source memory performance. (A) Proportion of “remember” and “familiar” responses to old and new items presented as hit rates, false alarm rates, and source memory rates are presented (with standard deviations) separately for “remember” and “familiar” responses. (B) Mean behavioral estimates of recollection and familiarity derived from the dual process model are presented (with standard deviations).

| (A) |  | Remember | Familiar |
| --- | --- | --- | --- |
| Object recognition performance | Hit rate | 0.47±0.14 | 0.32±0.08 |
|  | False alarm rate | 0.03±0.05 | 0.18±0.10 |
| Source memory accuracy |  | 0.47±0.13 | 0.33±0.07 |
| (B) |  | Recollection | Familiarity |
| Dual process estimate |  | 0.44±0.12 | 0.45±0.15 |

### ERP results

Before analyzing individual differences in ERP correlates of recognition, we conducted analyses to examine overall ERP old-new effects in order to be able to compare our results to previous reports of ERP differences between recollection and familiarity. We separately averaged ERPs for successfully recognized items associated with Remember responses, for recognized items associated with Familiar responses, and for Correct Rejection responses (Figure 2). These averages were done solely for visualization purposes. We report statistical comparisons between conditions in the following section. Averaged ERPs revealed a sustained negative deflection for all trials types, beginning approximately at 220 and lasting until around 400 ms, (Figure 2a-b) and a positive deflection from approximately 500 to 700ms after stimulus onset (Figure 2c-d).

**Figure 2.**
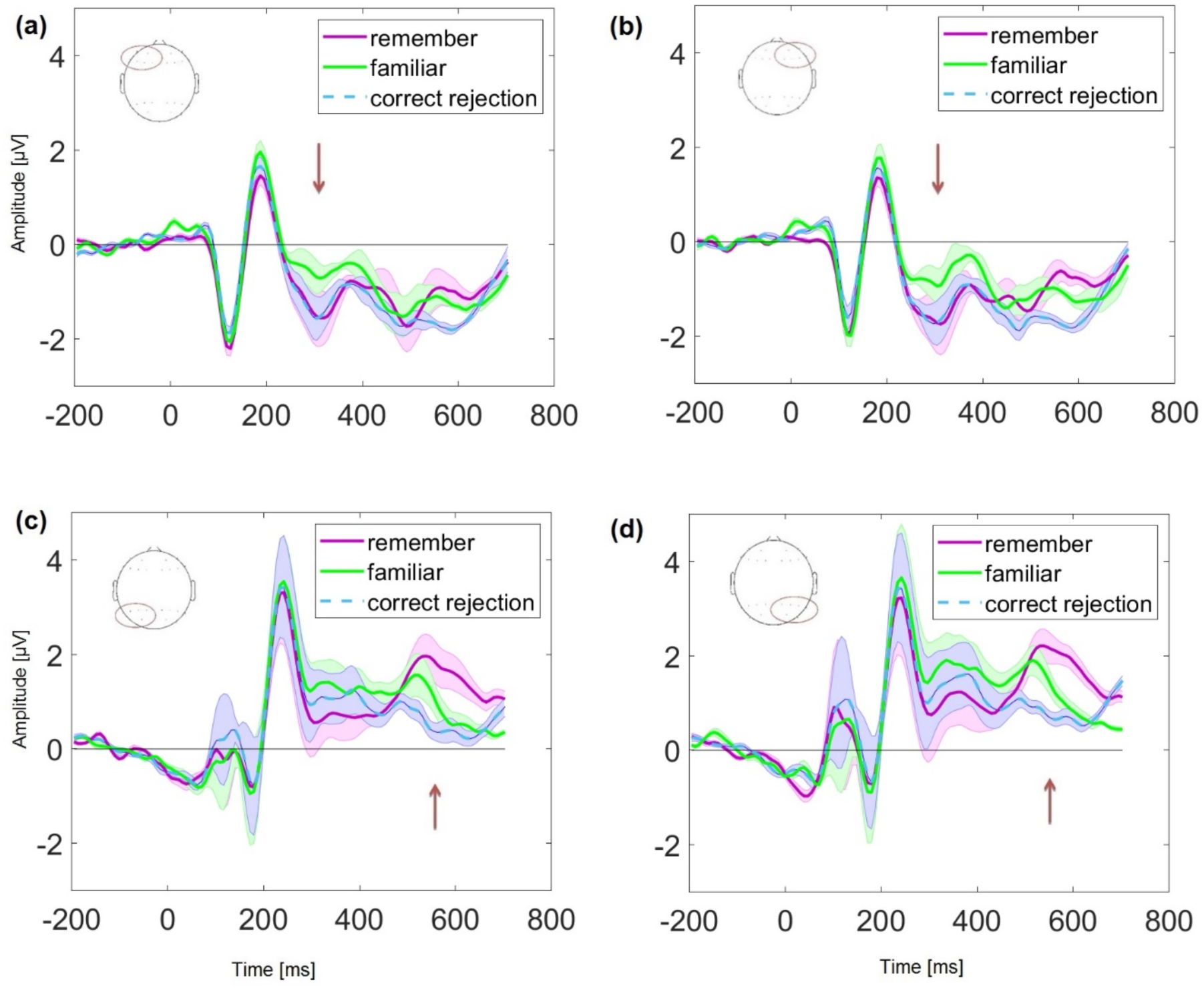
ERP correlates of recollection and familiarity. Grand averaged ERPs on Remember (purple), Familiar (green) and Correct Rejection (dashed blue) trials are separately averaged for four groups of channels split by frontal and parietal for each hemisphere (Woodruff et al., 2006). Arrows are meant to delineate time periods of interest, but do not indicate statistical comparisons: (a) left frontal (F1, F3, F5, F7, AF3, AF7), (b) right frontal (F2, F4, F6, F8, AF4, AF8), (c) left parietal (P1, P3, P5, P7, PO3, PO7), and (d) right parietal (P2, P4, P6, P8, PO4, PO8). Shaded areas represent standard deviation of the mean. Note that these average traces from electrode groups are presented for visualization purposes, but electrodes were analyzed separately in the data-driven statistical analyses.

In addition, to better visualize the familiarity and recollection ERP effects, we also present difference waveforms: Familiar-minus-Correct Rejection and Remember-minus-Familiar (Supplemental figure 3).

**Figure 3.**
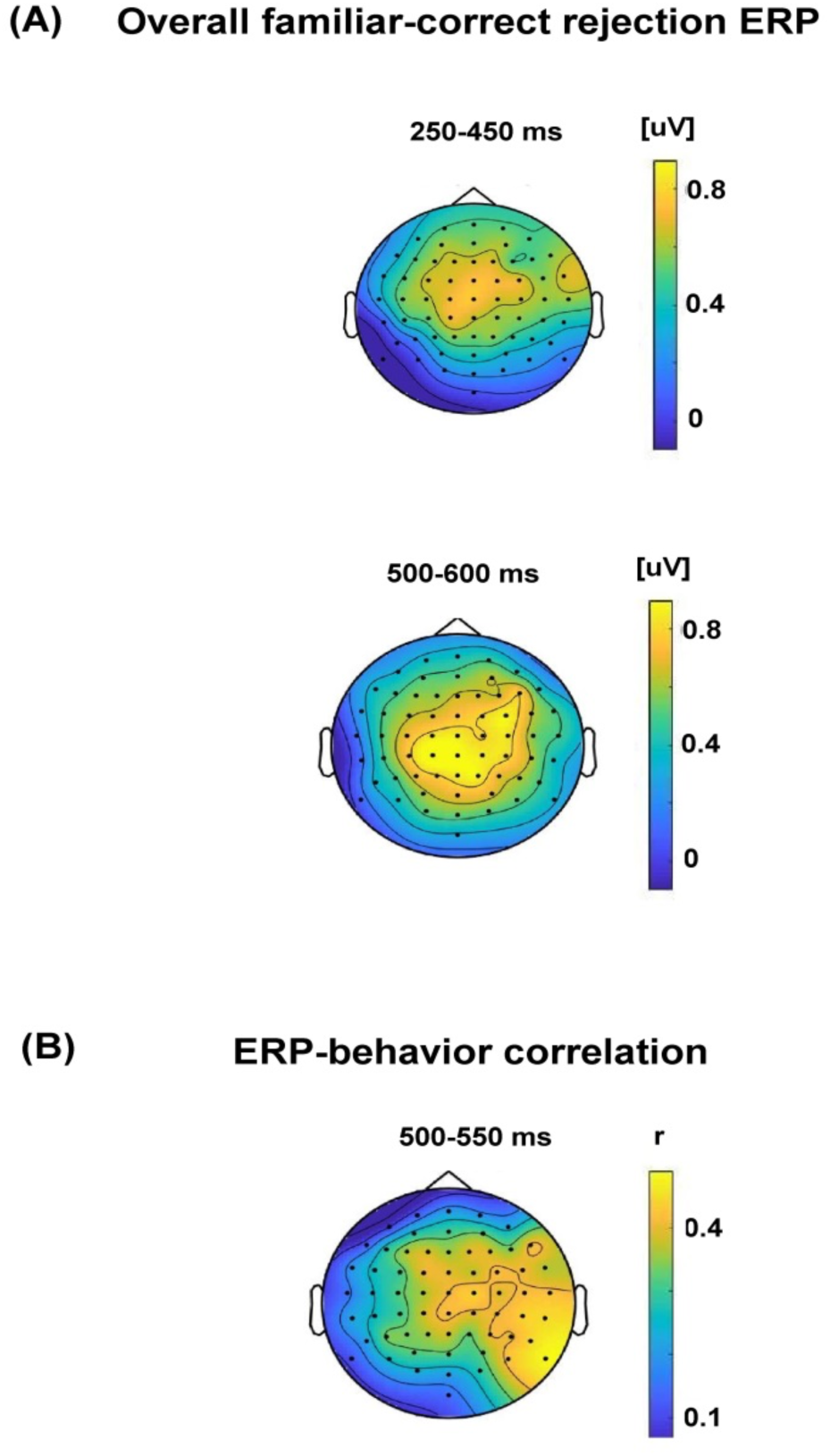
ERP correlates of familiarity. (A) Topographic maps illustrate distributions of mean ERP differences between familiar and correct rejection trials corresponding roughly to the significant clusters identified in the data-driven permutation analysis. (B) A topographic map illustrates a distribution of correlations between familiar - correct rejection ERP differences with dual process estimates of familiarity. Color bars show ERP voltage difference (panel A) or Pearson’s *r* correlation coefficient values (panel B).

To quantify ERP correlates of familiarity at the group level, we contrasted ERPs associated with familiar responses against ERPs associated with correct rejection responses. As described in the *Methods*, these, and all subsequent ERP analyses, were done using a data-driven non-parametric cluster-based permutation analysis procedure (Maris and Oostenveld, 2007). This method allows us to identify statistically significant differences between conditions, though it does not permit specific conclusions about the precise temporal or spatial extent of these differences (Sassenhagen and Draschkow, 2019). We can, however, identify the cluster extent in time and location as descriptive information about the observed data. As such, this analysis identified two spatiotemporal clusters that corresponded to the significant difference (*p*=0.001 cluster corrected for both clusters) in the observed data: from approximately 250 to 450 ms at frontal and fronto-central scalp sites, and from approximately 500 to 600 ms over central, centro-parietal, and right fronto-central sites. Figure 3a shows topographic distributions of the ERP differences between familiar and correct rejection trials corresponding to these time windows.

Next, we conducted data-driven analyses to identify ERP correlates of individual differences in familiarity-based recognition. This analysis revealed a significant cluster in the observed data (*p*=0.001 cluster corrected), extending approximately from 500 to 550 ms (Figure 3b). This correlation was most pronounced over right central and centroparietal areas. For completeness, we also analyzed correlations between familiar – correct rejection ERP differences and behavioral estimates of recollection. These analyses revealed no significant clusters. To summarize, we found that ERPs were sensitive to familiarity-based recognition, both at the overall group level and at the level of individual differences.

Next, to quantify ERP correlates of recollection, we contrasted ERPs associated with remember hits against ERPs associated with familiar hits. Analyses at the group level revealed significant differences corresponding to two spatiotemporal clusters (*p*=0.001 cluster corrected for both clusters). The first cluster extended from approximately 250 to 450 ms during which ERPs for remember trials were more negative than ERPs for familiar trials and had widespread scalp topography (Figure 4a upper panel). The second cluster extended from approximately 550 to 700 ms, and manifested as an enhanced positivity for remember trials compared to familiar trials. This latter effect had a centro-parietal scalp topography, largely consistent with prior reports of recollection-related ERP effects (Addante et al., 2012b; Curran, 2000; Duarte et al., 2004; Ranganath and Paller, 2000; Rugg et al., 1998b, 1998a; Rugg and Curran, 2007; Wilding, 2000; Wilding and Ranganath, 2012). Figure 4a shows topographic distributions of the ERP differences between remember and familiar trials during these time windows.

**Figure 4.**
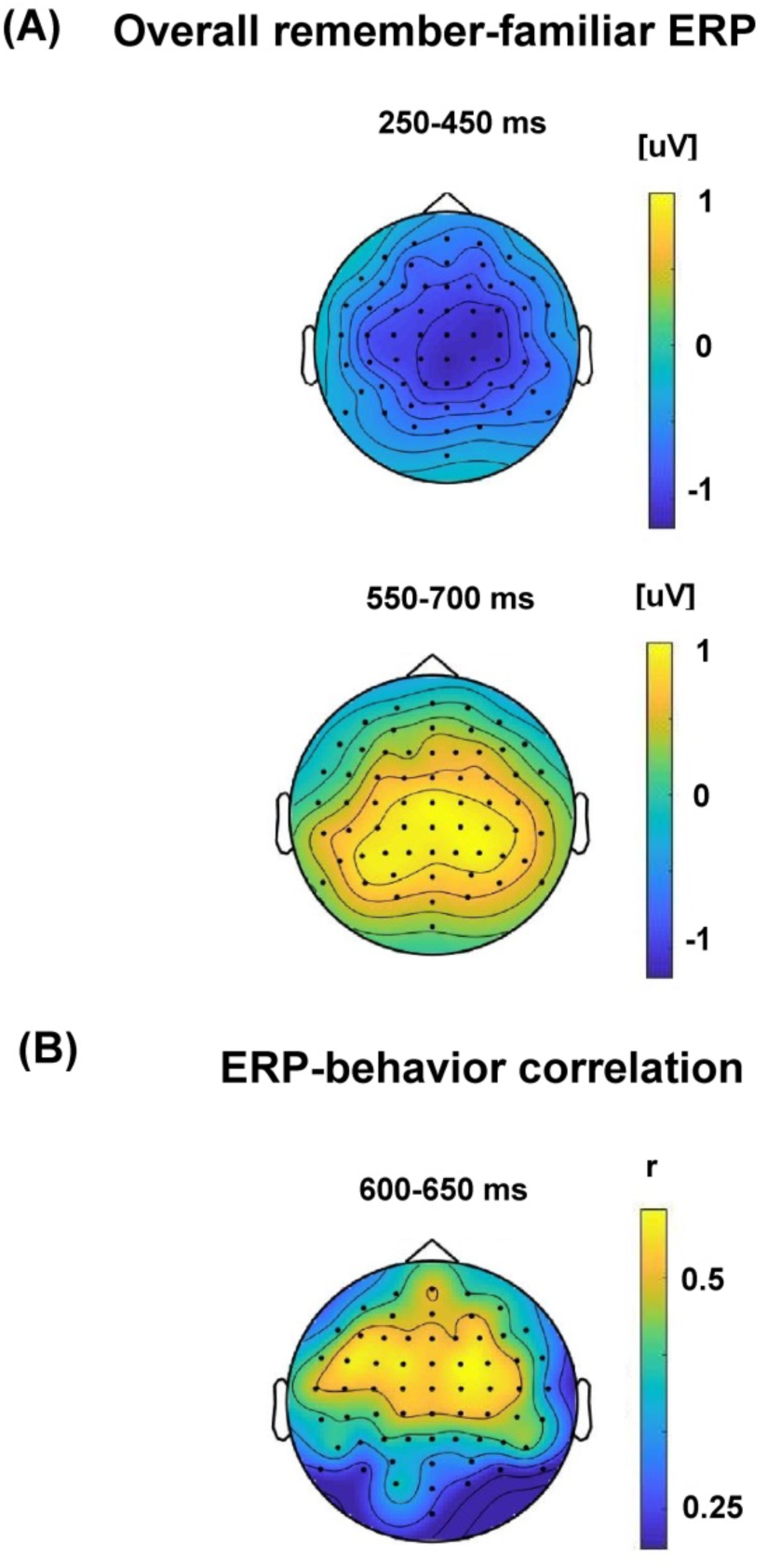
ERP Correlates of recollection. (A) Topographic maps illustrate distributions of mean ERP differences between remember and familiar trials during time windows that correspond roughly to the significant clusters identified in the data-driven permutation analysis. (B) A topographic map illustrates a distribution of correlations between Remember – Familiar ERP differences with dual process estimates of recollection. Color bars show ERP voltage difference (A) or Pearson’s *r* correlation coefficient values (B).

Having established significant remember – familiar differences at the group level, we next conducted a data-driven analysis to determine whether ERP differences between these trial types were positively correlated with dual process estimates of recollection. This analysis revealed a significant cluster (*p*=0.001 cluster corrected) in the observed data extending from 600 to 650 ms with a broad scalp distribution, particularly over central and fronto-central scalp sites (Figure 4b), where ERP amplitudes were positively correlated with recollection estimates. For completeness, we also analyzed correlations between remember – familiar ERP differences and behavioral estimates of familiarity. These analyses revealed no significant clusters. To summarize, these analyses revealed significant ERP correlates of recollection at the group level and at the level of individual differences.

To exclude a possible explanation of the results hinging on signal-to-noise ratio differences as a function of the number of trials contributing to the ERPs, we reran the analysis after equating the number of trials, which resulted in obtaining the same pattern of results. In supplemental figures 4 and 5 we also present topographic maps illustrating correlates of familiarity and recollection for the whole analyzed time window with highlighted electrode clusters on the basis of which the null hypothesis was rejected.

## Discussion

This study was designed to test the hypothesis that the neural correlates of recollection and familiarity-based recognition are predictive of individual differences in episodic memory performance. Data-driven analyses of ERPs during memory retrieval revealed overall effects broadly consistent with previous ERP studies of recognition memory, with an early mid-frontal ERP modulation that was enhanced for familiarity-based recognition, and a late posterior ERP modulation that was enhanced for recollection-based recognition. Critically, our data-driven analyses only revealed significant relationships between Familiar– Correct Rejection ERPs and individual familiarity estimates (Figure 3b), whereas we only found significant relationships between Remember – Familiar ERPs and individual recollection estimates (Figure 4b). ERP-behavior correlations were seen at a relatively late latency (>500ms post-stimulus) for both familiarity and recollection. These findings suggest that ERPs can provide useful markers of individual differences in recognition memory processes.

Although the central goal of the study was to look at individual differences in behavior and electrophysiology, we first wanted to determine the extent to which our results concurred with results from previous ERP paradigms. Consistent with a large body of evidence from ERP studies of recognition memory, we observed an early ERP old-new effect related to familiarity-based recognition and a late old-new effect related to recollection-based recognition. It is notable, however, that many other recognition memory correlates have been reported with time courses and scalp distributions that vary across paradigms (c.f., Wilding and Ranganath, 2012). This variance makes sense, because processes that support recognition memory can occur within 200 ms of the onset of a word or picture, and there is likely be extensive parallel, feedforward, and feedback processing throughout different brain networks (Clarke et al., 2011; Clarke and Tyler, 2014; Halgren et al., 2006; Marinkovic et al., 2003), resulting in field potentials that overlap in space and time at the scalp. Thus, the use of different types of stimuli (e.g., words, pictures, etc.) or different kinds of memory decision procedures across studies could likely engage different subprocesses that could impact the timing or topography of ERP correlates of memory (Bader et al., 2020; Busch et al., 2004; Taylor, 2002; Yonelinas, 2002).

In order to address this concern in an unbiased manner, we adopted a statistical technique, non-parametric cluster-based permutation testing (Maris and Oostenveld, 2007), that relies on the data to determine both significant time windows and electrode clusters. This was also adapted when identifying spatiotemporal clusters that correlate with behavioral memory measures. Our analyses revealed a rich picture, such that two different spatiotemporal clusters were associated with familiarity, and two different clusters were associated with recollection. Although our analysis methods do not permit precise inferences about the timing of these effects, it is notable that recollection and familiarity were each associated with clusters in relatively early and late time windows. This analysis enabled us to identify reliable effects without relying on assumptions from previous work, and, in turn, may have enabled us to uncover the relationships between electrophysiology and memory measures that have been previously mixed in other reports in the literature (Curran et al., 2006; Friedman and Johnson, 2000; Mecklinger, 2006, 2000; Olichney et al., 2000; Paller and Kutas, 1992; Rhodes and Donaldson, 2007; Rugg et al., 1998a; Smith, 1993; Tsivilis et al., 2001; Wilding and Ranganath, 2012). For instance, we identified different neural correlates of familiarity and recollection, but these results did not conform to the expectation (Curran et al., 2006) that familiarity-related neural processes should always precede those related to recollection.

A second key finding from this study is that ERPs also tracked individual differences in familiarity- and recollection-based recognition. Again, the use of data-driven approaches revealed results that might not have been obtained by assuming that individual differences in behavior should correlate with the magnitude of well-known ERP old-new effects. As shown in Figure 3, ERP correlations with familiarity estimates were seen over right posterior sites approximately 500-550 ms post-stimulus, a window which overlapped with the time window during which a significant group-level ERP familiarity effect was observed over central sites. Likewise, as shown in Figure 4, ERP correlations with recollection estimates were seen approximately 600-650 ms at frontal scalp sites, a time window that overlapped with a significant group-level ERP recollection effect with a centro-parietal topography.

One way to think about these results is that those who had higher behavioral estimates of recollection or familiarity showed ERP effects that were larger in magnitude than those who had lower recollection or familiarity estimates. To explore this possibility, we calculated grand averaged ERPs on remember, familiar and correct rejection trials for low and high (median split) familiarity estimate performers (Supplemental Figure 1) and low and high (median split) recollection estimate performers (Supplemental Figure 2). Both the familiarity (familiar – correct rejection difference within 500-600 ms time window) and recollection effects (remember – familiar difference within approximately 600-700 ms time window) were more pronounced in high than in low performers. Alternatively, it is possible that group-level ERP recollection and familiarity effects overlapped in time from separate ERP components that differentiated between those with high versus low performers. Although the latter possibility is less parsimonious, we cannot conclusively differentiate between topographic changes driven by increases in the strength of activity in the same configuration of neural sources vs. topographic changes driven by the involvement of different neural sources (Urbach and Kutas, 2002).

Our analyses were guided by models which propose that recollection and familiarity independently contribute to successful recognition memory. These models align with a vast body of evidence from lesion, intracranial EEG, and functional neuroimaging evidence demonstrating that familiarity disproportionately depends on representations of item-related information by the perirhinal cortex, whereas recollection disproportionately depends on binding of item and context information by the hippocampus (Davachi, 2006; Eichenbaum et al., 2007; Ranganath, 2010b; Ranganath and Ritchey, 2012). Other researchers, however, have been more agnostic about memory *content*, instead focusing on the idea that all retrieved information is summed together to provide an overall sense of the *strength* of a memory (Kelley and Wixted, 2001; Wixted, 2007). According to this view, “remember” and “familiar” responses reflect different points along a single one-dimensional continuum of memory strength (Kelley and Wixted, 2001). It is important to note that such single-process models do not attempt to characterize memory per se, but rather to account for the way decisions are made on a memory task—for instance, it is possible that there are qualitatively different neural signals for different kinds of memory content, and that the information is integrated into a single strength of evidence signal when making a behavioral response (Gold and Shadlen, 2007, 2001).

Our study was not designed to conclusively adjudicate between single- and dual-process models, but it is not clear that a single memory strength process would be sufficient to fully account for our results. If one were to assume that remember and familiar responses vary along a single memory strength continuum, and if ERPs reflect an aggregated measure of memory strength, then we would expect any ERP old-new effect to be larger for remember responses than for familiar responses. However, as we can see from the raw traces in Figure 2, prior to approximately 500 ms, there is an enhanced positivity for familiar trials that is virtually absent for remember trials. This might seem counterintuitive, but it aligns with the dual process model. According to models that assume independent contributions of recollection and familiarity to recognition (Aggleton and Brown, 1999; Atkinson and Juola, 1973, 1974; Atkinson et al., 1974; Eichenbaum et al., 1994; Jacoby, 1984, 1991, 1983; Jacoby et al., 1992; Jacoby and Dallas, 1981; Mandler, 1980; Norman and O’Reilly, 2003; Yonelinas, 2002, 2001a, 2001b, 1999, 1997, 1994), a familiar response is made only when familiarity is very high <u>and</u> recollection has failed. A remember response, in turn, happens when recollection is successful, even if the item’s familiarity is relatively low. Thus, the model would predict that an ERP correlate of familiarity can be very large on familiar trials and attenuated, or even absent, on remember trials (see also Diana et al., 2011). Moreover, if we solely consider behavioral performance, we can look at the associated, or source, information that can be retrieved when an item is successfully remembered. Although there is evidence that both familiarity and recollection can support accurate source memory (Addante et al., 2012b; Diana et al., 2011, 2008; Yonelinas, 2001a), recollection-based responses are more closely associated with retrieval of contextual details (Diana et al., 2012; Dimsdale-Zucker et al., 2018; Ranganath, 2010b, 2010a; Ranganath and Rainer, 2003). We observed significantly above chance source memory performance for items correctly given both familiar and remember responses. However, source memory performance on familiar trials was significantly lower than for remember trials. This fits with the dual-process account of recognition memory phenomenology (Park and Donaldson, 2019; Yonelinas, 2002, 1994; Yonelinas et al., 2010). Another key point is that recollection dual process estimates only correlated with remember – familiar ERPs and familiarity estimates only with familiar – correct rejection ERPs. Moreover, neither correlation effects overlapped in time and topography. The findings are compatible with a dual-process account. However, we acknowledge that the present non-parametric cluster-based permutation test methods do not allow us to make strong conclusions about the topography and timing of the familiarity and recollection ERP effects.

Another controversy in prior ERP studies of recognition memory has focused on the functional significance of the mid-frontal ERP old-new effects (Bridger et al., 2012; Paller et al., 2012, 2007; Voss et al., 2012; Voss and Federmeier, 2011; Voss and Paller, 2006; Yovel and Paller, 2004). Results from many studies have supported the idea that this old-new effect is enhanced during familiarity-based recognition, and that it is relatively insensitive to factors that influence recollection (Addante et al., 2012b; Bridger et al., 2012; Curran, 2004, 2000; Friedman and Johnson, 2000; Park and Donaldson, 2019; Rugg et al., 1998a; Rugg and Curran, 2007; Wilding and Herron, 2006; Wilding and Ranganath, 2012). However, a number of findings also support the idea that the mid-frontal old-new effect could instead reflect conceptual priming, which refers to more fluent processing of conceptual information that has been recently encountered (Guillem et al., 2001; Jelicic, 1995; Levy et al., 2004; Mitchell and Bruss, 2003; Nessler et al., 2005; Olichney et al., 2000; Paller et al., 2012, 2007; Ullsperger et al., 2000; Voss and Paller, 2007, 2006). For instance, thinking about the meaning of the word “banana” might make it more likely to come into mind when asked to generate the names of fruit words. Yovel and Paller (2004) used photographs of faces never seen before the experiment as stimuli, to isolate a pure familiarity effect. The authors found no association between familiarity and N400s and suggested that familiarity with faces may arise by a subset of the neural processing responsible for recollection, while the N400 reductions observed in the literature may reflect verbally mediated conceptual priming effects instead of familiarity. Another study (MacKenzie and Donaldson, 2007) obtained similar posterior old/new effect indexing familiarity for faces. However, in contrast to Yovel and Paller, the old/new effects associated with familiarity and recollection were topographically dissociable, consistent with a dual process view of recognition memory.

MacLeod and Donaldson (2017) also investigated the functional utility of the left parietal old/new effect using verbal stimuli. Their results revealed that ERP measures (defined as the mean ERP old/new difference within 500-800 ms post-stimulus averaged across left parietal electrodes: P1, P3, and P5) of retrieval were not related to behavioral performance. The authors concluded that the relationship between the left parietal effect and recollection is more complex than previously thought in the sense that the variation in the magnitude of the left parietal old/new ERP effect does not always reliably predict variation in episodic recollection between participants. However, the paper does not fully address the relation between ERPs and behavioral estimates of familiarity and recollection for several reasons: (1) the ERP effect was restricted only to the late left parietal effect, (2) the behavioral measures of recollection used in this study does not dissociate recollection from familiarity, and (3) only 20 participants were included in the correlation of R/K effect magnitude with behavioral data.

More recently, Wang et al. (2020) employed conceptually impoverished items (kaleidoscope images) as stimuli in a recognition memory test with a modified Remember/Know paradigm and they also observed that ERPs for Know hits were more positive than those for Correct Rejection items within 500-800 ms. Putting all these results together, there is considerable evidence suggests that the N400-like ERP effects are modulated by conceptual priming and familiarity (Bader and Mecklinger, 2017; Nessler et al., 2005; Wolk et al., 2004).

The controversy over ERP correlates of familiarity and conceptual priming relates to the broader question regarding the relationship between fluent processing and familiarity-based recognition memory. Substantial evidence exists to suggest that neural processes associated fluent processing of conceptual information are also related to familiarity—for instance, N400-like potentials occur in the perirhinal cortex during both conceptual priming and recognition memory (Nobre and McCarthy, 1995, 1994; Staresina et al., 2012).

Moreover, damage to the left perirhinal cortex impairs both conceptual priming and familiarity-based recognition memory (Bowles et al., 2007; Wang and Yonelinas, 2012a, 2012b; Wang et al., 2010). Finally, fMRI studies have shown that activity in the left perirhinal cortex during encoding predicts both conceptual priming and familiarity-based recognition, and perirhinal activity has been correlated with behavioral performance on both conceptual priming and familiarity-based recognition measures (Dew and Cabeza, 2013; Diana et al., 2010; Haskins et al., 2008; Heusser et al., 2013; Ranganath et al., 2004; Voss et al., 2009; Wang et al., 2015, 2014). Although these findings do not rule out the possibility that conceptual priming and familiarity can be dissociated (Paller et al., 2012), they are consistent with the broader idea that fluent processing of an item’s conceptual features can contribute to one’s subjective sense that the item is familiar (Mecklinger and Bader, 2020; Taylor and Henson, 2012; Wang and Yonelinas, 2012b).

In summary, the current study presents evidence to suggest that ERPs can be used to identify neural correlates of recollection and familiarity, both at the group level, and at the level of individual differences. The present findings provide support for the idea that ERPs can be used as biomarkers of underlying memory processes in healthy individuals, patient populations, or specific populations, like older adults or children (MacLeod and Donaldson, 2017). The combined use of behavioral and ERP measures, as in the present study, might be especially useful in the identification of those who are at risk for disorders such as Alzheimer’s disease (Xia et al., 2020). Additionally, the present results also highlight the potential value of data-driven analysis methods as a means to identify neural correlates of cognitive processes, complementing approaches that focus on specific, well-characterized components.

## Acknowledgement

This paper was supported by National Science Centre Poland under the project number 2018/02/X/ST6/02392 (KM) and a Vannevar Bush Faculty Fellowship from the Office of Naval Research Grant under the project number N00014-15-1-0033 (CR). At the time of final manuscript preparations, HDZ was supported by an NRSA F32 Fellowship (# F32EY032352). We thank Aria Fereydouni and Meyhaa Buvanesh for their help with data collection, Max Bluestone for sharing relevant EEG analysis code, and two anonymous reviewers for their comments on an initial submission of the manuscript.

## Supplementary materials

**Supplemental Figure 1.**
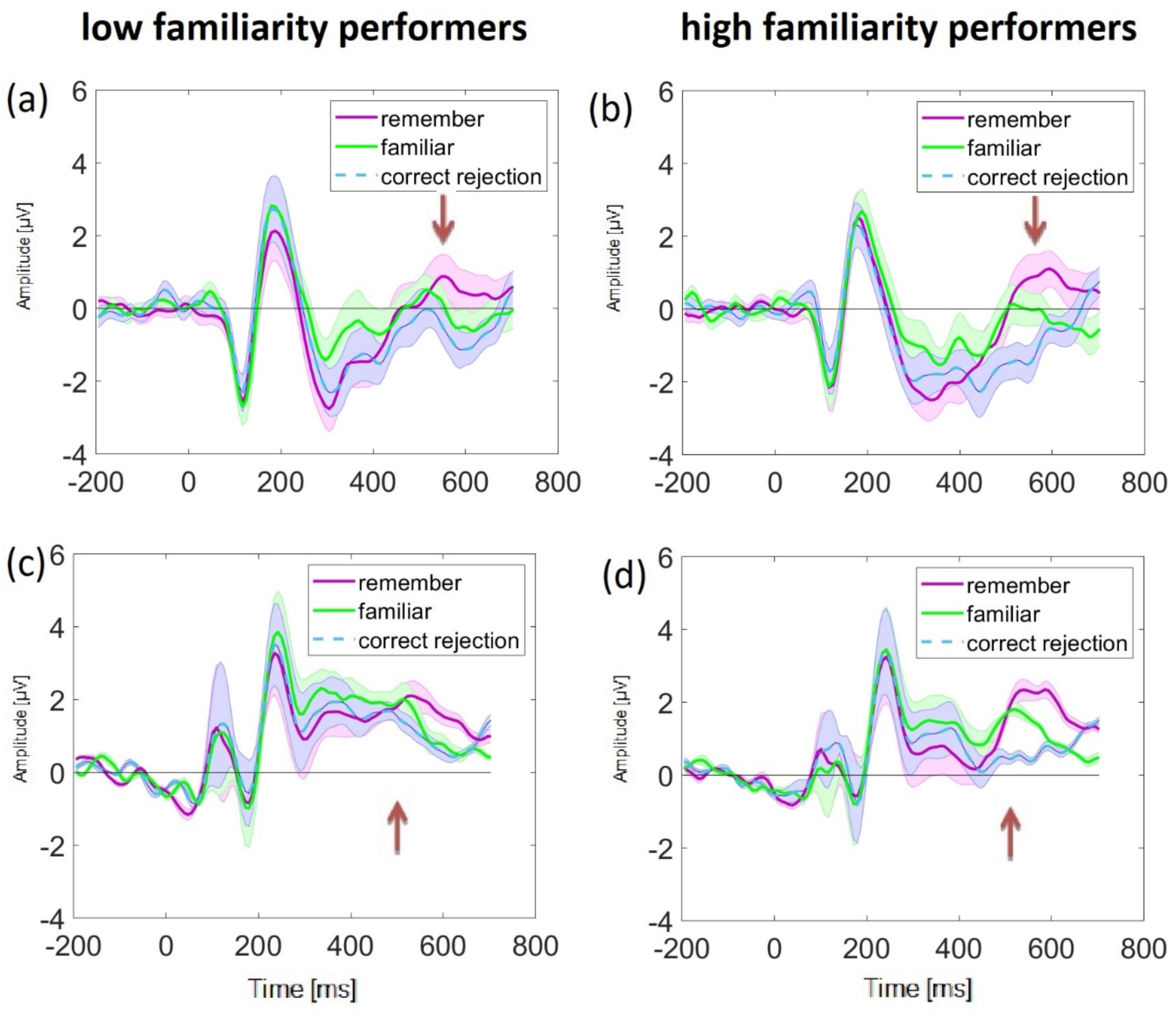
ERP correlates of familiarity. Grand averaged ERPs on Remember (purple), Familiar (green) and Correct Rejection (dashed blue) trials for low and high performers based on median split. Shaded areas represent standard deviation of the mean. Arrows are meant to delineate time periods of interest, but do not indicate statistical comparisons: (a) low familiarity estimate performers at Cz, (b) high familiarity estimate performers at Cz, (c) low familiarity estimate performers at right parietal (P2, P4, P6, P8, PO4, PO8), and (d) high familiarity estimate performers at right parietal (P2, P4, P6, P8, PO4, PO8). Note that the average traces from electrode groups are presented for visualization purposes, but electrodes were analyzed separately in the data-driven statistical analyses.

**Supplemental Figure 2.**
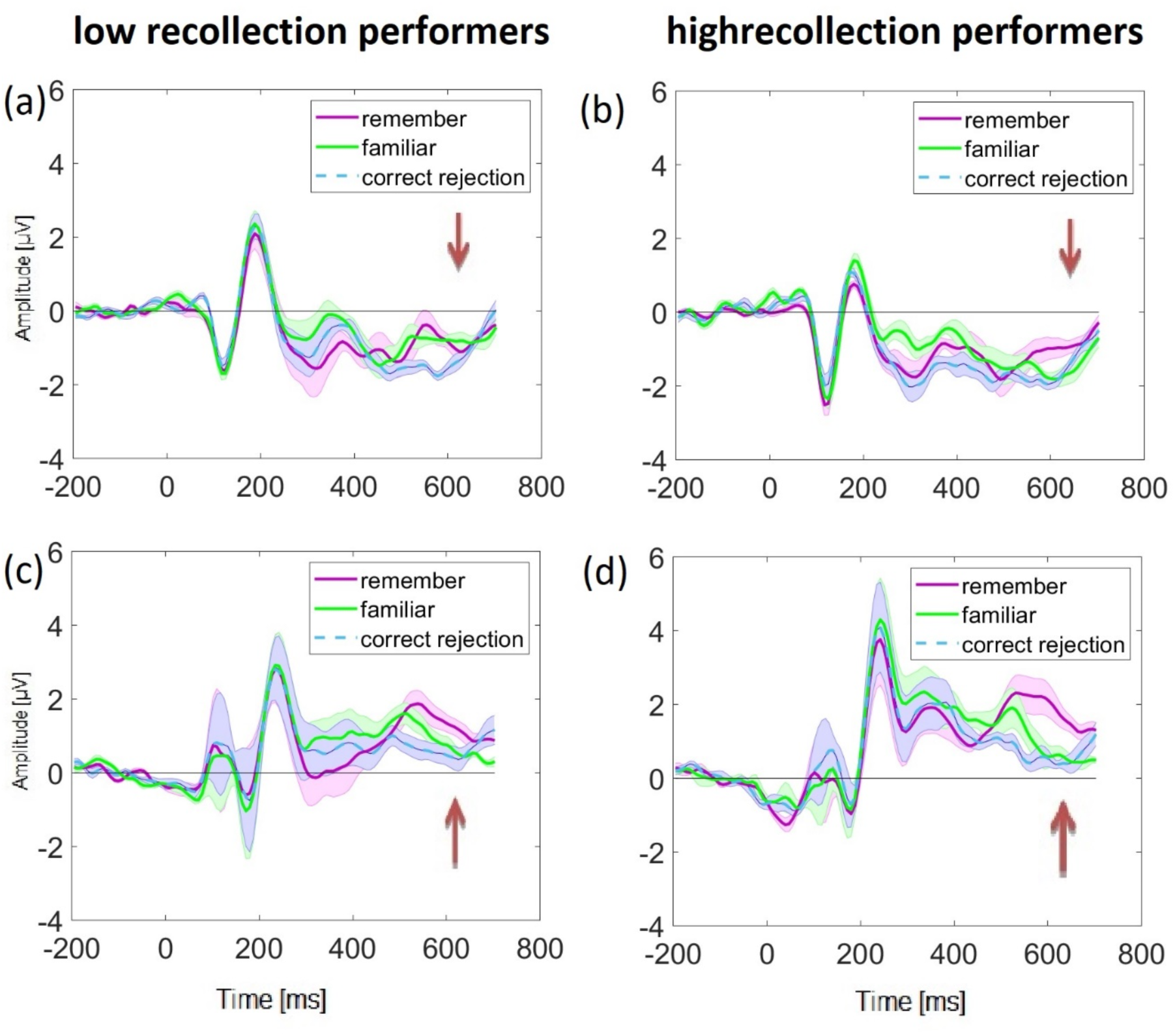
ERP correlates of recollection. Grand averaged ERPs on Remember (purple), Familiar (green) and Correct Rejection (dashed blue) trials for low and high performers based on median split. Shaded areas represent standard deviation of the mean. Arrows are meant to delineate time periods of interest, but do not indicate statistical comparisons: (a) low recollection estimate performers at frontal (F1, F3, F5, F7, AF3, AF7, F2, F4, F6, F8, AF4, AF8), (b) high recollection estimate performers at frontal (F1, F3, F5, F7, AF3, AF7, F2, F4, F6, F8, AF4, AF8), (c) low recollection estimate performers at parietal (P1, P3, P5, P7, PO3, PO7, P2, P4, P6, P8, PO4, PO8), and (d) high recollection estimate performers at parietal (P1, P3, P5, P7, PO3, PO7, P2, P4, P6, P8, PO4, PO8). Note that the average traces from electrode groups are presented for visualization purposes, but electrodes were analyzed separately in the data-driven statistical analyses.

**Supplemental figure 3.**
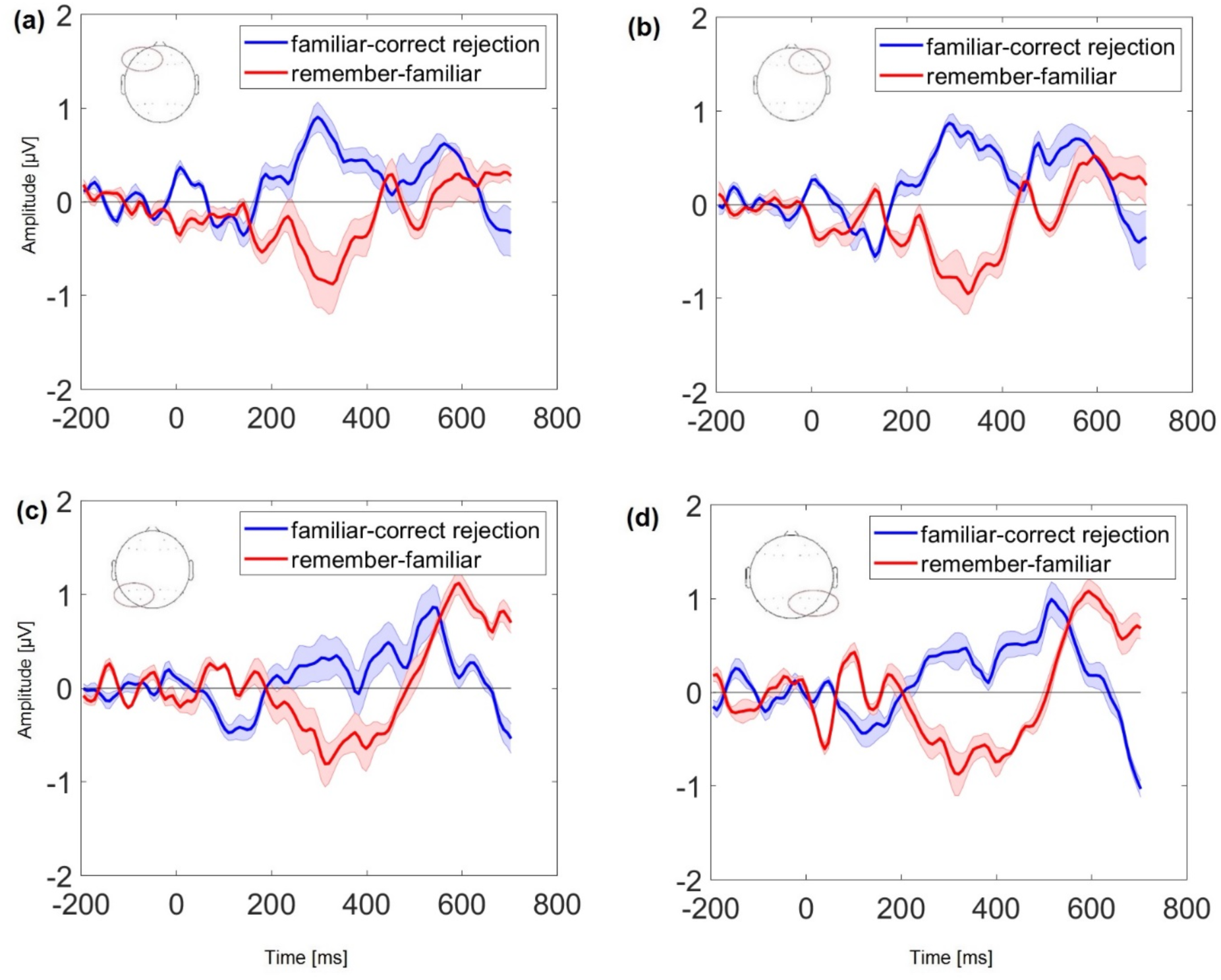
ERP correlates of recollection and familiarity. Grand averaged ERP difference waveforms: Familiar-minus-Correct Rejection (red) and Remember-minus- Familiar (blue), separately averaged for four groups of channels split by frontal and parietal for each hemisphere (Woodruff et al., 2006): (a) left frontal (F1, F3, F5, F7, AF3, AF7), (b) right frontal (F2, F4, F6, F8, AF4, AF8), (c) left parietal (P1, P3, P5, P7, PO3, PO7), and (d) right parietal (P2, P4, P6, P8, PO4, PO8). Shaded areas represent standard deviation of the mean. Note that these average traces from electrode groups are presented for visualization purposes, but electrodes were analyzed separately in the data-driven statistical analyses.

**Supplemental figure 4.**
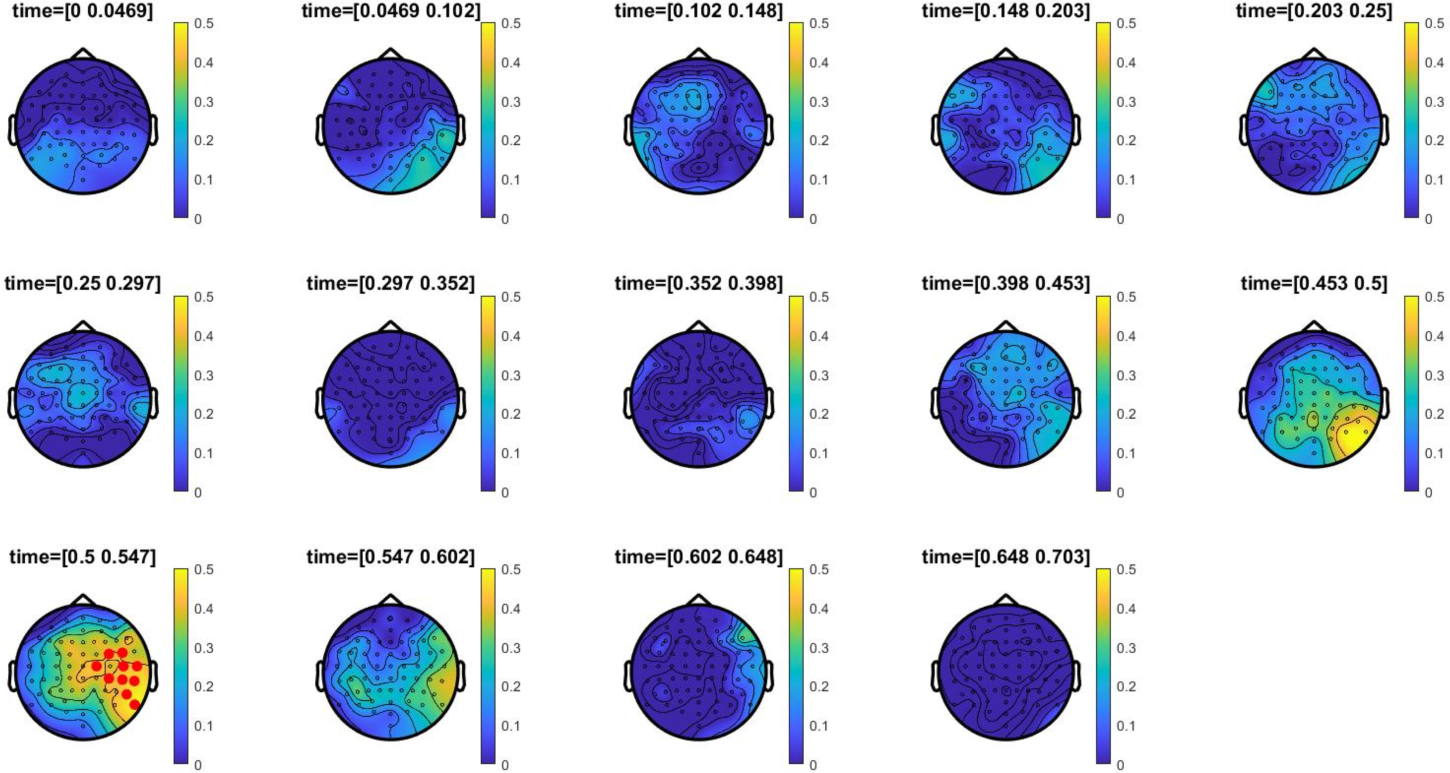
ERP correlates of familiarity. Topographic maps illustrate a distribution of correlations between Familiar - Correct Rejection ERP differences with dual process estimates of familiarity for 50 ms time bins. Electrode clusters on the basis of which the null hypothesis was rejected are highlighted with red asterisks. All timepoints and all 64 electrodes were included in the permutation test within specified 0-700 ms time window, at *P* < 0.05, cluster corrected. Color bars show Pearson’s *r* correlation coefficient values.

**Supplemental figure 5.**
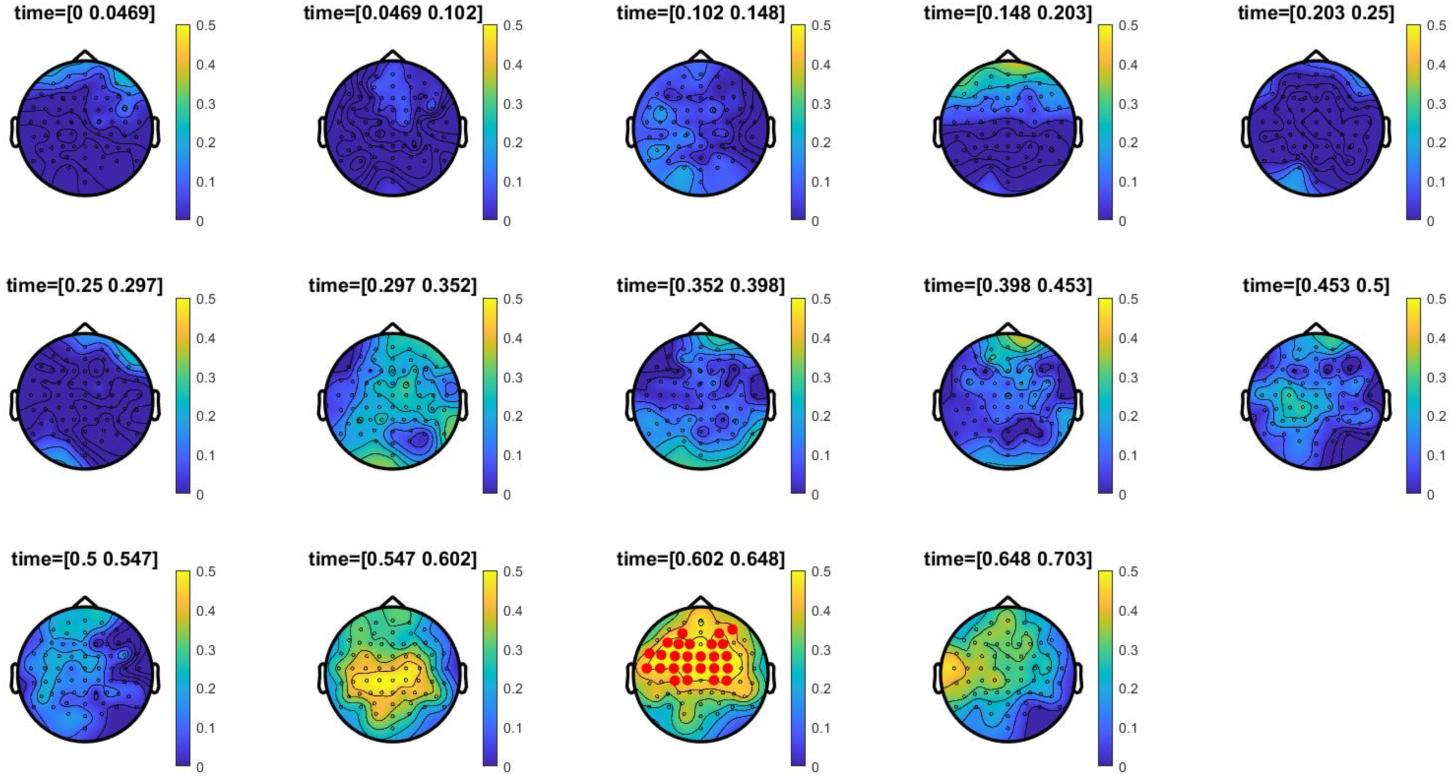
ERP Correlates of recollection. Topographic maps illustrate a distribution of correlations between Remember – Familiar ERP differences with dual process estimates of recollection for 50 ms time bins. Electrode clusters on the basis of which the null hypothesis was rejected are highlighted with red asterisks. All timepoints and all 64 electrodes were included in the permutation test within specified 0-700 ms time window, at *P* < 0.05, cluster corrected. Color bars show Pearson’s *r* correlation coefficient values (B).

**Supplemental Table 1.** Mean and standard deviation counts of confidence level by scored item response.

| Item response<br>(scored) | Highly<br>confident | Moderately<br>confident | Somewhat<br>confident | Not at all<br>confident |
| --- | --- | --- | --- | --- |
| Recollect | 66.36 (27.18) | 20.91 (14.21) | 4.19 (3.25) | 1.33 (0.58) |
| Familiar | 19.06 (14.06) | 18.42 (6.33) | 16.97 (8.84) | 3.76 (3.07) |
| Miss | 9.93 (9.18) | 12.53 (8.25) | 9.97 (5.88) | 2.89 (2.35) |
| Correct<br>rejection | 28.91 (19.59) | 24.39 (12.25) | 13.33 (7.88) | 4.11 (2.83) |

